# Loss of endogenous estrogen alters mitochondrial metabolism and muscle clock-related protein Rbm20 in female *mdx* mice

**DOI:** 10.1101/2024.02.08.579567

**Authors:** Cara A. Timpani, Didier Debrincat, Stephanie Kourakis, Rebecca Boyer, Luke E. Formosa, Joel R. Steele, Haijian Zhang, Ralf B. Schittenhelm, Aaron P. Russell, Emma Rybalka, Angus Lindsay

## Abstract

Female carriers of a *Duchenne muscular dystrophy* (*DMD*) gene mutation manifest exercise intolerance and metabolic anomalies that may be exacerbated following menopause due to the loss of estrogen, a known regulator of skeletal muscle function and metabolism. Here, we studied the impact of estrogen depletion (via ovariectomy) on exercise tolerance and muscle mitochondrial metabolism in female *mdx* mice and the potential of estrogen replacement therapy (using estradiol) to protect against functional and metabolic perturbations. We also investigated the effect of estrogen depletion, and replacement, on the skeletal muscle proteome through an untargeted proteomic approach with TMT-labelling. Our study confirms that loss of estrogen in female *mdx* mice reduces exercise capacity, tricarboxylic acid cycle intermediates and citrate synthase activity but that these deficits can be offset through estrogen replacement therapy. Furthermore, ovariectomy downregulated protein expression of RNA binding motif factor 20 (Rbm20), a critical regulator of sarcomeric and muscle homeostasis gene splicing, which impacted pathways involving ribosomal and mitochondrial translation. Estrogen replacement modulated Rbm20 protein expression and promoted metabolic processes and the upregulation of proteins involved in mitochondrial dynamics and metabolism. Our data suggests that estrogen mitigates dystrophinopathic features in female *mdx* mice and that estrogen replacement may be a potential therapy for post-menopausal DMD carriers.

## 1. Introduction

Metabolic and mitochondrial dysfunction is a well-established nuance of dystrophin-deficient skeletal muscle^1–6^. Mutation of the X-chromosome residing *DMD* gene that leads to the loss of the cytoskeletal dystrophin protein, causes Duchenne muscular dystrophy (DMD), a progressive and eventually fatal neuromuscular disease that predominantly affects males. Loss of control of several metabolic systems, e.g., mitochondrial tricarboxylic acid (TCA) cycle and electron transport chain (ETC) function, glycolysis, and purine nucleotide metabolism, perpetuates the DMD pathobiology by compromising cellular energy homeostasis and promoting oxidative stress^1^. However, the root cause of these metabolic disturbances is unclear. Some purported theories include (1) loss of nitric oxide synthesis due to deficient dystrophin-neuronal NO synthase (nNOS) binding^7^; (2) loss of calcium homeostasis owing to enhanced sarcolemmal fragility and calcium leak channel hyperactivity caused by dystrophin deficiency resulting in persistent induction of the mitochondrial permeability transition pore^8^; (3) dysregulated purine nucleotide cycle function leading to purine degradation and reduced ATP recovery potential^9^; (4) upregulated micro RNA (miR)-379, resulting in reduced mitochondrial ATP synthase function^10^; and (5) dysregulated mitophagy and mitobiogenesis signalling^11^. Mitochondrial and metabolic dysfunction is evident in dystrophic myoblasts^12^ and early in the pathological dystrophinopathy milieu leading to muscle degeneration^13^. Targeting mitochondria/metabolism can ameliorate disease pathology in the *mdx* mouse model of DMD, e.g., via mitochondria transplantation^14^ or targeted drug therapies (ginovistat targets HDACs^15^, urolithin A activates mitophagy^11^, dimethyl fumarate targets TCA anaplerosis^16^).

In addition to DMD patients and animal models that carry biallelic *DMD*/*Dmd* gene mutations, mitochondrial/metabolic dysfunction is evident in female carriers of a DMD gene mutation. Approximately two-thirds of DMD patients inherit mutations from carrier mothers (the other one-third of cases occur due to *de novo* mutation^17^). These carriers frequently report exercise intolerance, being unable to perform the same extent of muscle work for a matched workload as non-carriers^18,19^. They also take longer to recover high phosphate energy stores post-exercise^18,19^ and show sharp increases in serum CK^20^, suggesting susceptibility of muscle fibres to mechanical damage even though dystrophin is expressed (albeit sub-optimally^21^). However, rarely do they manifest progressive muscle wasting (<10% of cases)^22^. Female DMD carriers might be particularly susceptible to manifestations of DMD as they reach menopause and the protective effects of endogenous estrogen production subside. Indeed, several case studies highlight muscle wasting and pathology causing disability in ∼50-year-old DMD carriers^23,24^.

The *mdx* mouse model of DMD carries a point mutation in the *DMD* gene and shows various degrees of metabolic perturbation despite an overall milder phenotype than DMD patients. Both male and female mice carry the biallelic gene mutation due to inbreeding and share natural history homology except that females are more resistant to laboratory-based stressors (e.g., handling and scruffing^25,26^). To explore this aspect further, we recently assessed the phenotype in ovariectomised female *mdx* mice to test the hypothesis that estrogen was protective against heightened stress responses^25,26^. Although estrogen depletion did not exacerbate the muscle-specific phenotype, our serum metabolomics screen indicated that mitochondrial TCA cycle metabolite levels were reduced but were effectively replenished by estradiol (E2) therapy^26^. The protective effects of estrogen on skeletal and cardiac muscle have been demonstrated previously^27^ but never linked to metabolic dysfunction in the context of dystrophin deficiency. Pharmacological estrogen receptor modulation has been pursued as a potential therapy for DMD. The selective estrogen receptor alpha (ERα) agonist, tamoxifen, was initially shown in *mdx* mice to improve whole-body strength, skeletal muscle function and reduce cardiac pathology^28^. More recent work demonstrates it lessens contractile dysfunction in stem-cell-derived cardiomyocytes^29^. However, clinical trials in Switzerland could not show that tamoxifen modified the natural history of DMD progression despite improvements in some functional measures^30,31^. The potential of ERα modulation may not be fully realised in male patients who express fewer ER’s than female counterparts, but it could be useful to protect female carriers from progressive disease in the peri-/menopausal years.

The aim of this study was to explore the impact of estrogen depletion on muscle mitochondrial metabolism and the potential for E2 replacement therapy to protect against metabolic maladaptation in female *mdx* mice. We studied the effect of estrogen manipulation on the skeletal muscle proteome using an untargeted proteomic approach with TMT-labelling and demonstrated that estrogen levels modulate the expression of RNA binding motif factor 20 (Rbm20). Since Rbm20 is regulated by the muscle molecular clock program^32^, our data suggest that estrogen depletion alters mitochondrial metabolism potentially through modulating the muscle circadian rhythm.

## 2. Materials and Methods

### 2.1 Study Approval

This research was approved by the Deakin University Animal Ethics Committee (G09-2020). Mice used in this study were housed in accordance with the Deakin University Animal Welfare Committee standards and cared for in accordance with the Australian Code for Care and Use of Animals for Scientific Purposes.

### 2.2 Experimental Design & Animals

Muscles analysed in this study were collected from mice utilised in Lindsay *et al.* 2021 and 2023^25,26^. Briefly, 8-week-old C57BL/10ScSn-*Dmd^mdx^*/Arc female mice subjected to a sham surgery, ovariectomy (OVX) or OVX+E2 supplementation and were purchased from ARC (Western Australia, Australia; *n* = 15/group). Mice underwent surgery in which the ovary and oviduct was exteriorised and either returned into the peritoneal cavity (Sham) or cauterised between the uterine horn and oviduct (OVX and OVX+E2). Once the muscle wall was closed, either a placebo (Sham and OVX) or E2 pellet (OVX+E2; 3.4 µg/day E2 release) was inserted under the skin. Mice recovered for two weeks prior to shipment to Deakin University. All mice were housed in groups of four/cage on a 12/12 h light/dark cycle with food and water provided *ad libitum*. Mice acclimatised for one week before undergoing scruff restraint. A forced downhill treadmill exercise-to-fatigue test was performed where the treadmill was set at 0 m/min for two min before increasing to 10 m/min for 1 min. The treadmill speed increased by 1 m/min until a speed of 15m/min was reached and then maintained for 15 min. A second scruff restraint was performed one week later and 15 min after the final scruff restraint, mice were sacrificed via CO_2_ asphyxiation and cervical dislocation. Blood was collected, and muscles were harvested, snap-frozen, and stored at -80°C until required.

### 2.3 Metabolomics

Metabolomics was performed in Lindsay and Russell 2023^25^. Briefly, serum samples were extracted using a 1:1 acetonitrile/methanol solution, vortexed, incubated at 4°C for 10 min and centrifuged at 4°C for 10 min on maximum speed. The supernatant was transferred into a HPLC insert and the polar metabolites were separated and analysed via mass spectrometry at Metabolomics Australia (Bio21 Molecular Science and Biotechnology Institute, University of Melbourne, Parkville, Australia).

### 2.4 Muscle oxidative capacity

OCT-covered gastrocnemius, tibialis anterior (TA) and diaphragm muscles were cryo-sectioned (10 µm at -15°C) and succinate dehydrogenase (SDH) capacity was quantified as described by us previously^16,33^.

### 2.5 Mitochondrial enzyme activity

Activity of citrate synthase (CS), an enzyme of the TCA cycle, was measured spectrophotometrically in gastrocnemius homogenates as described by us previously^16,33^. To complement assessment of SDH activity in muscle sections, activity of SDH/mitochondrial ETC complex II (CII) was assessed on isolated mitochondria from gastrocnemius muscles according to the manufacturer directions (Abcam, ab228560). Mitochondria were isolated as performed by us previously^34^.

### 2.6 Proteomics

#### 2.6.1 Protein extraction from samples and enzymatic digestion

Quadriceps were cryo-pulverised with subsequent solubilisation in 5% sodium dodecyl sulphate (SDS) 10 mM Tris HCl, heat inactivation performed at 95°C for 10 min followed by probe sonication 3 x 30 s rounds and centrifuged at 13,000 g for 5 min to clarify. The supernatant of each sample was transferred to a new tube, and protein concentration was measured using a BCA kit (Thermo Fisher, #23225) according to the manufacturer’s instructions. Samples were then processed using the S-trap protocol as per the manufacturer’s instructions (Protifi^35^). Normalised amounts of protein were reduced and alkylated with 10 mM TCEP (Thermo, #77720) and 40 mM chloroacetamide (Sigma, C0267-100G) with incubation at 55 °C for 15 min. Enzymatic digestion was performed using Trypsin (Promega, V528X) at a 1:50 wt:wt ratio alongside Lys-C at a 1:25 wt:wt ratio (Promega, VA1170) at 37 °C for 16 h. Digestion efficiency was greater than 94% for this analysis.

#### 2.6.2 Tandem Mass Tag (TMT) labelling and fractionation

Each sample was labelled with the TMTpro 18plex reagent set (Lot:XJ351218 and XK350589, Thermo Scientific) according to the manufacturer’s instructions utilising a singular reference channel (126) containing all samples pooled. Individual labelled samples were then pooled into plexes, and high-pH RP-HPLC was used to generate 36 fractions concatenated to 12, which have been acquired individually by LC-MS/MS to maximize identifications. Labelling efficiency was determined to be greater than 97%.

#### 2.6.3 Liquid Chromatography Mass Spectrometry Protocol

Liquid chromatography-mass spectrometric (LC-MS) analysis was conducted using an□Orbitrap Eclipse Tribrid mass spectrometer (Thermo Scientific, Breman, Germany) and Nano LC system (Dionex Ultimate 3000 RSLCnano). The samples were loaded onto in Acclaim PepMap 100 trap column (100 μm x 2 cm, nanoViper, C18, 5 μm, 100Å; Thermo Scientific) and separated on an Acclaim PepMap RSLC (75 μm x 50 cm, nanoViper, C18, 2 μm, 100Å; Thermo Scientific)□analytical column. The peptides were resolved by increasing concentrations of buffer B (80% acetonitrile/0.1% formic acid) and analyzed via 2 kV nano-electrospray ionisation. The mass spectrometer operated in data-dependent acquisition mode using in-house optimized parameters with 120 min of chromatographic separation used for each fraction. Briefly, the acquisition used three FAIMS compensation voltages (-40, -55, -70 V) operated under standard resolution with an iron transfer tube temperature of 300□ with a carrier gas flow rate of 4.6 L/min. Precursor ion scans were performed at a 120,000 resolution from 400 - 1,600 m/z, an AGC target of 250% and ion injection time set to auto. Peptide fragmentation and reporter tag quantification were performed synchronously (10 per duty cycle per compensation voltage) with the fragmentation spectra generated in the ion trap using CID with turbo scan rate; MS3 reporter ion measurements were performed in the orbitrap with a resolution of 50,000. A precursor isolation filter was used for the selection of ions with a 0.7 Da and 50% envelope fit. Dynamic exclusion was applied for 60 s across all compensation voltages with only one charge state per precursor selected for fragmentation. In addition, the use of real-time searching was performed with a close out of 10 peptides per protein within each injected fraction; this utilised the human SwissProt proteome with 1% false discovery filtering applied during acquisition.

#### 2.6.4 Mass spectrometric data analysis

The raw data files were analyzed using Sequest within Proteome Discoverer (v2.5.0.400, Thermo Scientific)□to obtain protein identifications and their respective reporter ion intensities using in-house standard parameters. The Mouse SwissProt proteome containing only reviewed sequences (accessed June 2023) was used for protein identification at a 1% false discovery rate (FDR) alongside a common contaminants database. Reporter ion quantifiers used a unique plus razor with analysis centred on protein groups for shared peptide sequences using all peptides for abundance determination; quantitative values were corrected for stable isotope label impurities according to the manufacturer’s values.

#### 2.6.5 Bioinformatic data analysis

Protein level data was exported and analysed utilising TMT-Analyst, the latest addition to the Monash Proteomic Analyst Suite (https://analyst-suites.org/), which is built upon the foundations of LFQ-Analyst^36,37^. Briefly, prior to normalisation, proteomic data was filtered for high-confidence proteins from already determined master proteins representing groups. Proteins marked as contaminants and proteins not quantified consistently (condition N/2 + 1) across the experiment were removed. The remaining missing values were imputed using the missing-not-at-random (MNAR) method, assuming the missingness was due to low expression for such proteins, which were then normalised using the variance-stabilising-normalisation (VSN) method. Both imputations and VSN were conducted by the DEP package^38^. The limma package^39^ from R Bioconductor was used to generate a list of differentially expressed proteins for each pair-wise comparison. A cut-off of the adjusted *p*-value of 0.05 (Benjamini-Hochberg method) and a log2 fold change of 0.75 was applied to determine significantly regulated proteins in the different pairwise comparisons. We utilised the protein abundance data obtained from the LFQ-Analyst suite and conducted differential pathway analysis using Correlation Adjusted MEan RAnk gene set test (CAMERA)-intensity based analysis within Reactome (v83)^40^. Briefly, this method uses limma-based differential analysis on mean protein abundances within pathways across samples with adjusted FDR correction to control for multiple hypothesis testing. Using this approach enables the detection of alterations in pathways that will likely remain unobserved when examining individual differentially expressed proteins. This is because individual proteins might not provide comprehensive insights into how a specific pathway is changing when assessed through global, untargeted proteomic-level statistical comparisons. The FDR of <0.05 is reported.

### 2.7 Statistics

Data are reported as mean ± SEM unless stated otherwise. Normal distribution of the data was assessed using a Shapiro-Wilk test and all data passed the normality test except for the forced downhill treadmill exercise-to-fatigue test. This data was analysed using a Kruskall-Wallis test and a Dunn’s multiple comparison test was performed post hoc. All other data, excluding proteomics (see section 2.6.5), were analysed using one-way ANOVA with Tukey’s post hoc test on GraphPad Prism version 10.0.3 (GraphPad Software, Boston, Massachusetts, USA). α was set at 0.05 and trends are reported at <0.1.

## 3. Results

### 3.1 Estrogen replacement therapy recovers exercise capacity and mitochondrial metabolism

The impact of estrogen depletion on exercise capacity of female *mdx* mice was assessed via a forced downhill treadmill exercise-to-fatigue test. OVX reduced the exercise capacity of female *mdx* mice by ∼64% compared to Sham (Figure 1A; *p*<0.0001) with E2 replacement normalising exercise capacity to Sham levels (∼133% increase, Figure 1A; *p*<0.01). Since exercise capacity is intricately linked to mitochondrial metabolism, the metabolomics data set was analysed to assess TCA cycle intermediates. Metabolomic analysis captured six of nine TCA cycle intermediates, and when pooled, OVX reduced TCA metabolite content compared to Sham (Figure 1B; *p*<0.0001). This effect was mostly accounted for by significantly reduced succinate levels (Figure 1C; *p*<0.0001). As with exercise capacity, E2 replacement therapy restored the TCA metabolite pool (Figure 1B; *p*<0.05), especially succinate concentration (Figure 1C; *p*<0.05).

**Figure 1.**
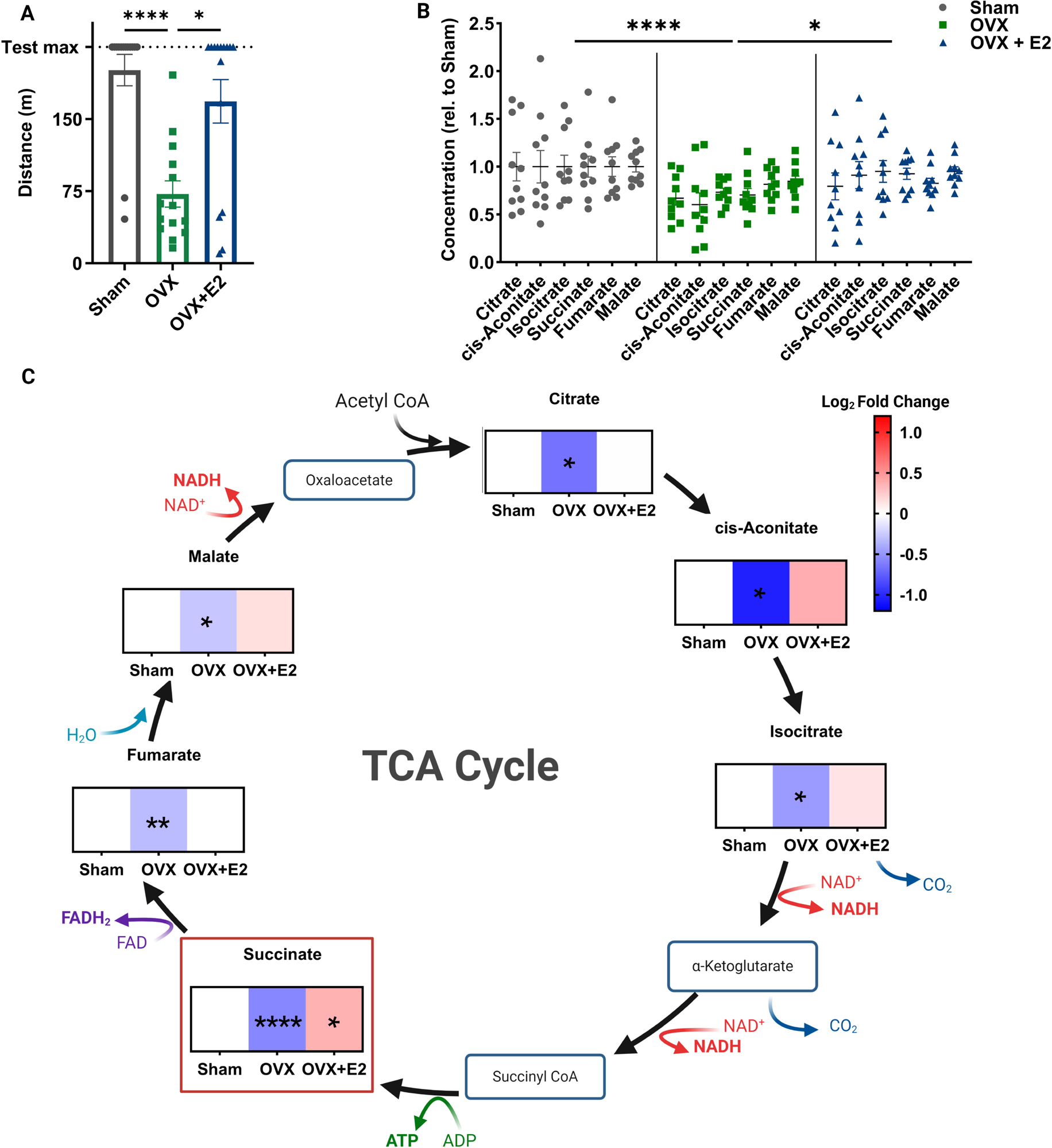
Estrogen depletion reduces exercise capacity and mitochondrial metabolism intermediates in female *mdx* mice that can be recovered with estradiol (E2) replacement. (A) Exercise capacity, as assessed by a forced downhill treadmill exercise-to-fatigue test, was reduced in ovariectomised (OVX) mice compared to Sham with E2 replacement improving exercise capacity in OVX mice. (B) Concentration of pooled tricarboxylic acid (TCA) cycle intermediates is reduced following OVX but is restored after E2 replacement therapy. (C) The reduction in pooled TCA cycle intermediates was driven by a significant decrease in succinate concentration which was recovered with E2 replacement. Data in (C) is expressed as the log_2_ fold change. \**p*<0.05, \*\**p*<0.01, \*\*\**p*<0.001, \*\*\*\**p*<0.0001. Sham *n*=10-14; OVX *n*=10-14; OVX+E2 *n*=10-15.

### 3.2 Estrogen replacement recovers mitochondrial content in gastrocnemius but reduces SDH capacity in TA and diaphragm

Since OVX affected mitochondria-specific metabolic intermediates, the activity of key mitochondrial enzymes was determined, including (1) CS, the pace setter of TCA cycle flux and a well-established biomarker of mitochondrial content, and (2) SDH, which couples TCA cycle flux to ETC function. Since metabolic intermediates were quantified in serum, it was necessary to confirm alterations in muscle mitochondrial enzyme activity were associated with the serum metabolic signature as was shown previously^41^. CS activity of whole muscle homogenates was reduced in OVX compared to Sham gastrocnemius (Figure 2A; *p*<0.01) and was recovered via E2 replacement (Figure 2A; *p*<0.0001), indicative of either increased substrate flux or mitochondrial density. Despite an ∼40% reduction in mean values, there was no statistically significant effect of OVX on SDH activity of isolated mitochondria from the gastrocnemius (Figure 2B; *p*=0.6491). There was, however, a trend for E2 replacement to increase SDH activity compared to OVX mice (Figure 2B; *p*=0.0898). SDH capacity of the gastrocnemius was also assessed through histological staining to provide information on potential fiber type shifts (Figure 2C; *p*>0.05). SDH capacity of two other muscles – one being the TA, which is commonly analyzed for histology, and the diaphragm, which is typically impacted as the dystrophic disease worsens, were also assessed. Interestingly, OVX significantly increased the SDH capacity of TA (Figure 2D; *p*<0.01) and trended to do the same in the diaphragm (Figure 2E; *p*=0.0691). In both muscles, E2 replacement normalized SDH capacity to Sham levels (Figure 2D & E respectively; *p*<0.01). While the proportion of less, more, and highly oxidative fibers did not shift in the gastrocnemius (Figure 2F & Supp Figure 1A^I-III^; *p*>0.05), OVX drove a less oxidative phenotype in the fast-twitch fiber predominant TA (Figure 2G & Supp Figure; *p*<0.01), which was normalized by E2 replacement (Figure 2G & Supp Figure 1B^I-III^; *p*<0.01). In contrast, OVX increased the proportion of more oxidative fibers in the diaphragm compared to Sham (Figure 2H & Supp Figure 1C^II^; *p*<0.05) while E2 replacement tended to increase the proportion of highly oxidative fibers in OVX diaphragm (Figure 2H & Supp Figure 1C^III^; *p*=0.086).

**Figure 2.**
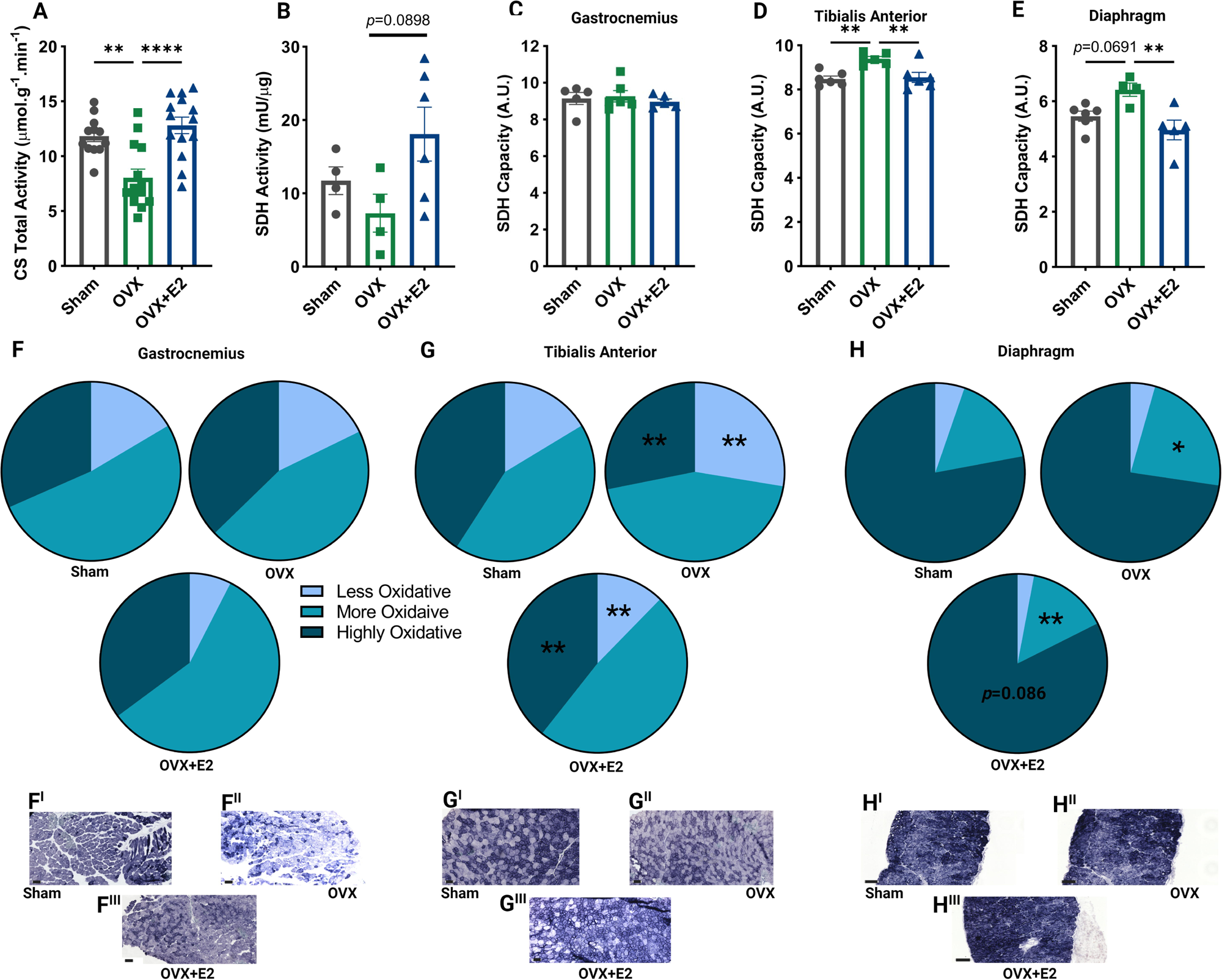
Estrogen depletion reduces citrate synthase (CS) but not succinate dehydrogenase (SDH) activity in gastrocnemius of female *mdx* mice. (A) CS activity, a surrogate for mitochondrial content, was reduced in ovariectomized (OVX) gastrocnemius compared to Sham, which was recovered with estradiol (E2) replacement. (B) While SDH enzyme activity was comparable between Sham and OVX mitochondria isolated from gastrocnemius, there was a trend for E2 replacement to increase SDH activity (*p*=0.0898). Using SDH histological staining of muscle sections, SDH capacity of gastrocnemius was comparable across all groups (Sham, OVX and OVX+E2; C), but was increased in tibialis anterior (D), with a trend detected in diaphragm (*p*=0.0691; E), following OVX. In both tibialis anterior and diaphragm, E2 replacement decreased the SDH capacity (D & E). (F-G) The proportion of less, more, and highly oxidative fibers is shown for gastrocnemius, tibialis anterior and diaphragm with representative images (F-H^I-III^). \**p*<0.05, \*\**p*<0.01, \*\*\*\**p*<0.0001. Sham *n*=5-12; OVX *n*=4-14; OVX+E2 *n*=5-14. Scale bar= 100µm.

### 3.3 Estrogen depletion downregulates the protein expression of Rbm20, which is normalized by estrogen replacement and upregulates metabolic proteins

E2 replacement recovered both exercise capacity and CS activity. Therefore, the effect of OVX, and subsequently E2 replacement, on proteins involved in metabolism were investigated. To address this with an unbiased method, we performed TMT labelling of muscle lysates following trypsin and Lys-C digestion and subjected these to LC-MS analysis. More than 5,400 distinct proteins were identified by untargeted, label-based proteomics using TMT and interestingly, only one protein, Rbm20, was found to be significantly regulated using a log_2_ fold change cutoff of 0.75 and an adjusted *p* value of <0.05. Rbm20 (accession Q3UQS8), which regulates post-transcriptional splicing of sarcomeric and other muscle homeostasis genes, was downregulated in OVX compared to Sham muscle (Figure 3A & A^I^; *p*<0.0001). To understand whether OVX impacts families of proteins, which were not detected as significant due to the log_2_ fold change cutoff and *p* value criteria, pathway analysis (via Reactome) of the entire proteome dataset was conducted^42^. A total of 59 pathways were impacted by OVX (Figure 3B). Sixteen pathways were upregulated after OVX and particularly involved processes associated with vision (e.g., retinoid metabolism and transport, visual phototransduction, diseases associated with visual transduction, the canonical retinoid cycle in rods (twilight vision)). Of the 43 pathways that were downregulated following OVX, a significant proportion were associated with translational activity (e.g., eukaryotic translation initiation and elongation, cap-dependent translation initiation, mitochondrial translation). Restoration of estrogen through E2 replacement led to the significant upregulation of the Rbm20 protein (compared to OVX; Figure 4A and A^I^) as well as the modulation of a further 61 proteins, upregulating 29 and downregulating 32. Since Rbm20 was impacted by both OVX and E2 replacement, we probed the proteome dataset for its splicing targets^43^ to determine if the down- and upregulation of Rbm20 impacted subtle, but biologically relevant, changes on downstream proteins. Of the 30 proteins captured in our proteomics set, three proteins were downregulated following OVX (Camk2d and Pdlim3 (*p*<0.05) and Dtna (*p*<0.01); Figure 4B) and one protein was upregulated (Myh7, *p*<0.05). E2 replacement upregulated the expression of five proteins (Camk2d and Pdlim5 (*p*<0.05), Dtna (*p*<0.01) and Mlip and Sh3kbp1 (*p*<0.0001)) and downregulated the expression of two proteins (Nexn (*p*<0.05) and Mecp2 (*p*<0.001)).

**Figure 3.**
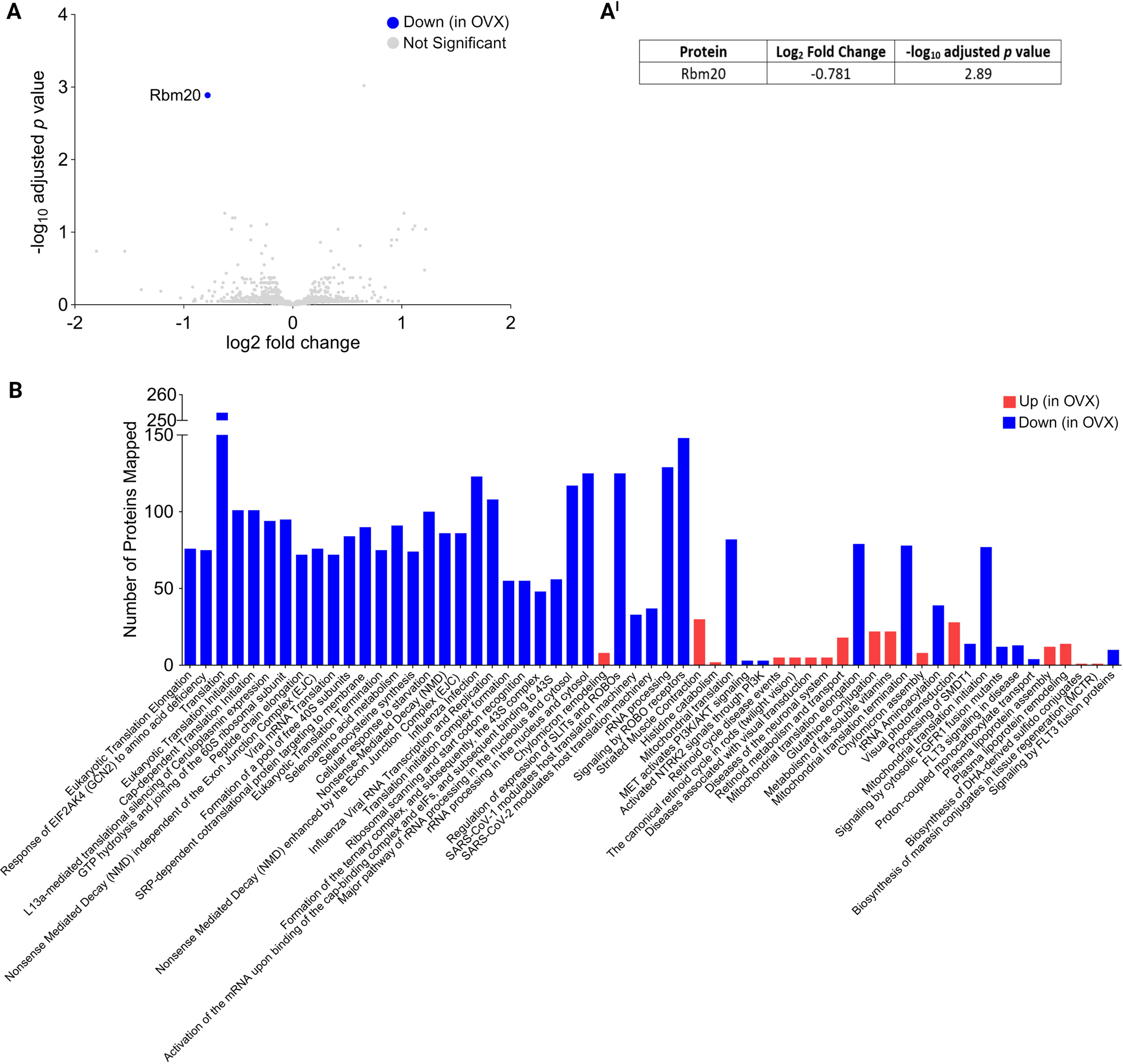
Estrogen depletion downregulates Rbm20 expression in female *mdx* mice and impacts 59 cellular processes. (A & A^I^) Of the ∼5,400 proteins identified through proteomics, only Rbm20 was impacted by ovariectomy (OVX) and downregulated in comparison to Sham. (B) Pathways analysis of the entire proteomics dataset identified that 59 pathways were impacted by OVX – 16 pathways were upregulated compared to Sham while 43 pathways were downregulated. Order of pathways is presented as most significantly impacted (i.e., eukaryotic translation elongation) to least significantly impacted (i.e., signaling by FLT3 fusion proteins). Sham *n*=5; OVX *n*=6.

**Figure 4.**
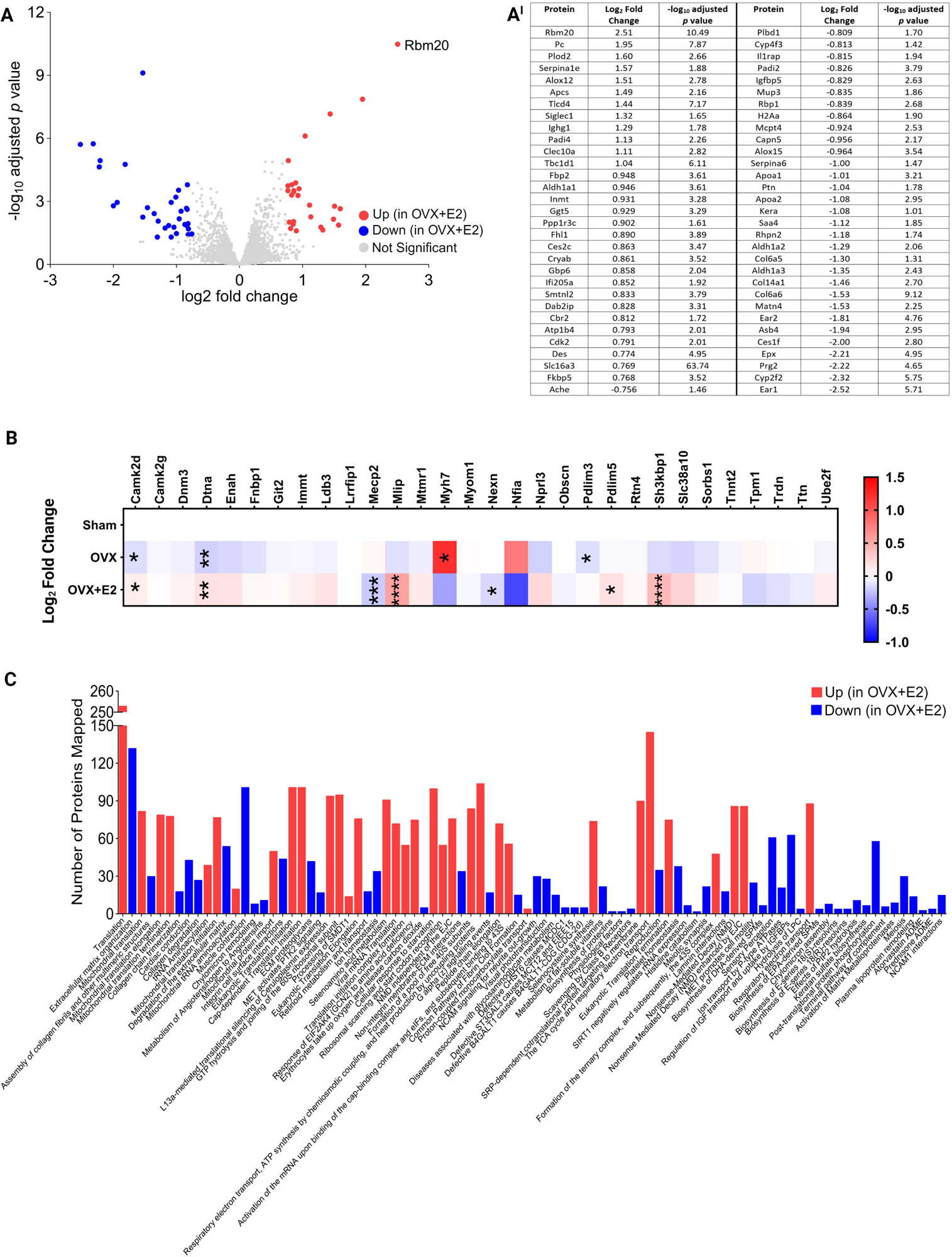
Estrogen replacement upregulates Rbm20 expression in female *mdx* mice and modulates the expression of a further 61 proteins. (A & A^I^) Rbm20 expression was upregulated following estradiol (E2) replacement as well as modulated expression of a further 61 others. (B) The impact of ovariectomy (OVX), and E2 replacement, is demonstrated on splicing targets of Rbm20. (C) Pathways analysis of the entire proteomics dataset identified that 88 pathways were impacted by E2 replacement – 34 pathways were upregulated compared to OVX muscle while 54 pathways were downregulated. Order of pathways is presented as most significantly impacted (i.e., translation) to least significantly impacted (i.e., NCAM1 interactions). \**p*<0.05, \*\**p*<0.01, \*\*\**p*<0.001, \*\*\*\**p*<0.0001 OVX v Sham or OVX+E2 v OVX. Sham *n*=5; OVX *n*=6; OVX+E2 *n*=6.

Of the 29 proteins that were upregulated by E2 replacement, a critical protein associated with orchestrating metabolic flux between the cytosolic glycolytic pathway and the mitochondrial TCA cycle, pyruvate carboxylase (Pc), was upregulated indicating a mechanism for the metabolomic shift. Indeed, pathways analysis revealed that of the 88 pathways modulated by E2 replacement (34 upregulated, 54 downregulated), key metabolic processes were upregulated, including the TCA cycle and respiratory ETC (Figure 4C). Furthermore, mitochondrial processes, including translation and import, were upregulated, suggesting an overall increase in mitochondrial activity. This motivated us to probe the proteomics dataset for key proteins driving these changes. Several proteins in the large and small subunit of the mitoribosome were downregulated by OVX (Mrpl12, Mrpl14, Mrpl20, Mrpl32, Mrpl50, Mrps28 and Mrps34, *p*<0.05, Figure 5A) with E2 replacement upregulating the expression of six large subunit proteins (*p*<0.05) and 17 small subunit proteins (*p*<0.05-0.0001). Markers of mitochondrial dynamics, including Cs, Mfn1 and Mfn2 (mitofusins), Opa1 (optic atrophy 1), Perm1 (PGC-1/ERR-induced regulator in muscle 1), Tfam (mitochondrial transcription factor A) and Vdac 1-3 (voltage-dependent anion channel), were not affected by OVX (Figure 5B; *p*>0.05) but Mfn1, Perm1, Tfam and Vdac2 (*p*<0.01) were significantly increased by E2 replacement. Notably, Cs protein expression was not affected.

**Figure 5.**
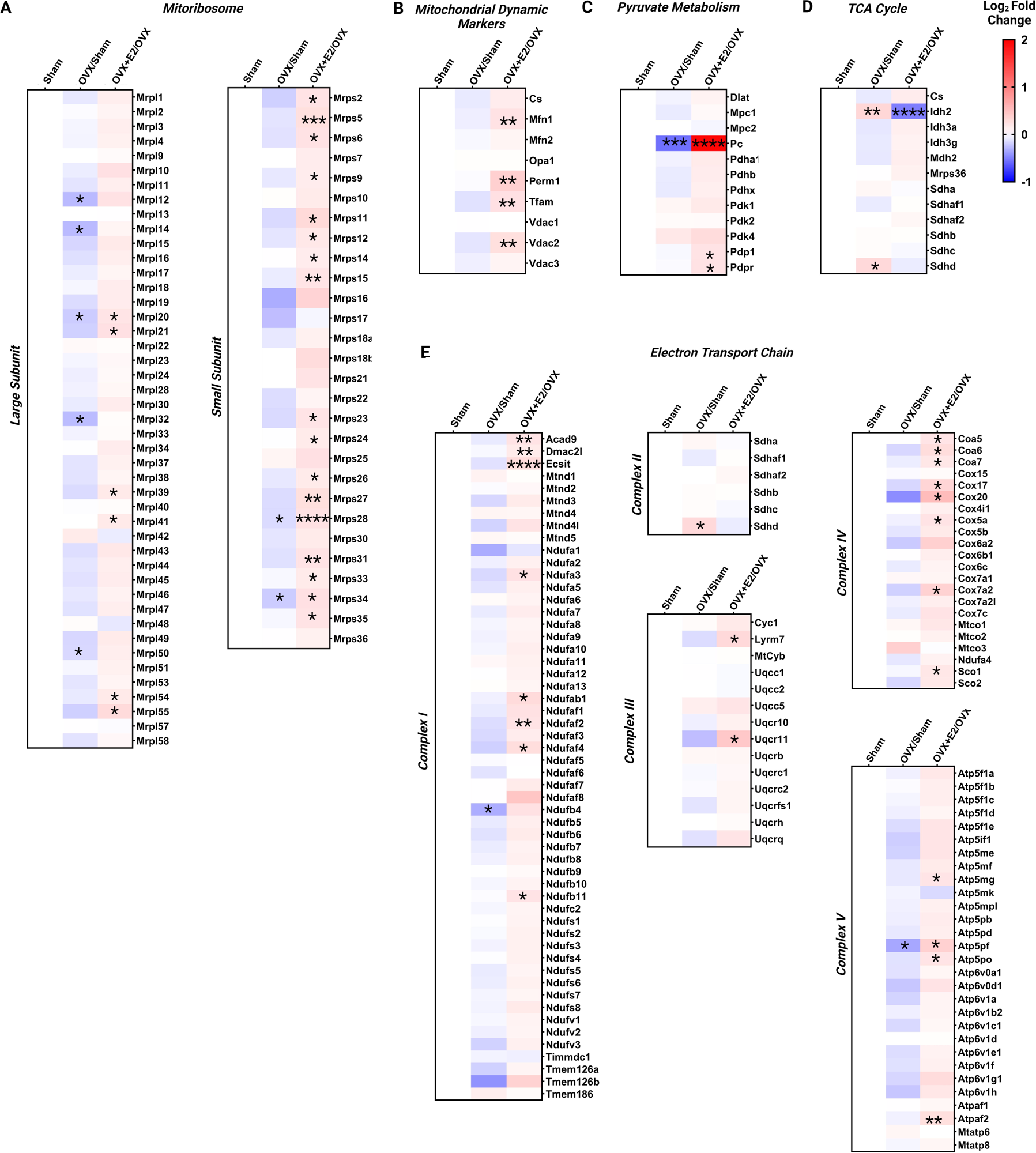
Estrogen replacement upregulates proteins associated with mitochondrial dynamics and metabolism in female *mdx* mice. The proteomics dataset was probed for proteins associated with the mitoribosome, mitochondrial dynamics and metabolism and are displayed as heatmaps with ovariectomized (OVX) compared to Sham and OVX+E2 (estradiol) compared to OVX. The impact of OVX, and E2 replacement, is demonstrated on (A) the mitoribosome, (B) markers of mitochondrial dynamics, (C) pyruvate metabolism, (D) the tricarboxylic acid (TCA) cycle and (E) the electron transport chain. \**p*<0.05, \*\**p*<0.01, \*\*\**p*<0.001, \*\*\*\**p*<0.0001 OVX v Sham or OVX+E2 v OVX. Sham *n*=5; OVX *n*=6; OVX+E2 *n*=6.

Since Pc was significantly upregulated following E2 replacement (Figure 4A and A^I^), proteins associated with pyruvate metabolism were explored. While it was not detected as a significantly dysregulated protein based on the log_2_ fold change criteria of 0.75 (Figure 3A), loss of estrogen downregulated the expression of Pc in OVX mice compared to Sham (Figure 5C; *p<*0.001). OVX had no impact on any of the other 11 proteins associated with pyruvate metabolism (*p*>0.05). However, E2 replacement upregulated expression of Pdp1 (pyruvate dehydrogenase phosphatase 1) and Pdpr (pyruvate dehydrogenase phosphatase regulatory subunit) compared to OVX mice (*p*<0.05). We next assessed proteins of the TCA cycle since we observed an increase in TCA cycle intermediates (Figure 1B-D) and CS activity (Figure 2A) following E2 replacement. Of the proteins captured in this dataset, OVX increased the expression of isocitrate dehydrogenase 2 (Idh2; Figure 5D; *p*<0.01) and a component of the SDH complex (Sdhd; *p*<0.05) compared to Sham. While E2 replacement was unable to modulate Sdhd expression (*p*>0.05), it reduced the expression of Idh2 (*p*<0.0001). Since pathway analysis revealed the ETC as an upregulated process, we next evaluated proteins associated with each respiratory complex (CI-V). Of the 54 CI structural subunits or assembly factors we could detect, only Ndufb4 was downregulated in OVX compared to Sham (Figure 5E; *p*<0.05). Eight proteins were upregulated in response to E2 replacement – Ndufa3, Ndufab1, Ndufaf4 and Ndufb11 (*p*<0.05) and Acad9 and Ndufaf2 (*p*<0.01). Only one of the six proteins associated with CII (Figure 5E) was affected by OVX (Sdhd; *p*<0.01), and this upregulation was not modified following E2 replacement (*p*>0.05). OVX did not affect any of the proteins associated with either CIII (14 proteins) or CIV (19 proteins; Figure 5E; *p*>0.05). However, E2 replacement significantly upregulated expression of the CIII proteins Lyrm7 and Uqcr11 and the CIV proteins Coa5-7 and Cox 5a, Cox7a2, Cox17, Cox20 and Sco1 (*p*<0.05). One of the 29 proteins associated with CV was downregulated following OVX (Atp5pf; Figure 5E; *p*<0.05) and E2 replacement normalized its expression (*p*<0.05). E2 replacement also upregulated the expression of the CV subunits Atp5mg, Atp5po (*p*<0.05) and Atpaf2 (*p*<0.01).

## 4. Discussion

Despite expressing dystrophin (albeit to varying degrees^44,45^), female DMD carriers manifest exercise intolerance and metabolic anomalies, including delayed recovery of high phosphate energy^18^. These are likely to be exacerbated following menopause since estrogen is known to regulate skeletal muscle function and metabolism by positively regulating mitochondrial biogenesis, respiratory chain respiration and lipid metabolism^46^. Our study confirms that OVX reduces exercise capacity, TCA cycle intermediate concentration and reduces CS activity in female *mdx* mice. Moreover, replacement of estrogen with E2 offset these deficits, supporting the idea that loss of estrogen could exacerbate dystropathology in female DMD carriers. In OVX mice, proteomics analysis identified a reduction in Rbm20, a critical regulator of sarcomeric and muscle homeostasis gene splicing. This was associated with changes in the skeletal muscle proteome that reflected reduced ribosomal and mitochondrial activity. Estradiol replacement attenuated the downregulation of the Rbm20 protein that was observed in the OVX-only group and promoted metabolic and mitochondrial processes, suggesting that estrogen allays the dystrophic phenotype of female DMD carriers.

Exercise intolerance is a well-documented effect following the loss of estrogen (previously reviewed in^46,47^) and our findings support this notion since OVX mice had a significantly reduced exercise capacity (∼64%), which was normalized by E2 replacement. Since exercise capacity is linked with metabolism, we probed our published serum metabolome dataset^26^ to elucidate a pathway/s which could, in part, explain the observed deficits in exercise. We determined that TCA cycle intermediates were reduced in OVX mice – specifically succinate concentration. In ER-related-α (ERRα) KO mice, muscle TCA cycle intermediates, including succinate, are reduced following exercise, however we did not see an accumulation of other intermediates (e.g., citrate or cis-aconitate) as observed in Perry *et al*^48^. This could be explained by either the difference in intensity of the exercise tests employed or the fact that our study used OVX to reduce estrogen levels while Perry *et al.* utilized ERRα KO to reduce estrogen binding to skeletal muscle while maintaining circulating levels. We did not quantify whether OVX completely abolished circulating estrogen (confirmed with reductions in uterus mass), thus our findings may be reflective of residual circulating estrogen, which has been previously demonstrated in rats one-month post OVX^49^. Irrespective, our data indicate that loss OVX impacts TCA cycle flux. For succinate, the reduced concentration may be reflective of increased succinate metabolism by a higher SDH capacity (significant in the TA and trend in the diaphragm). This is plausible as we observed an increase in the Sdhd subunit of CII, which is responsible for terminal electron transfer flow to CoQ and coordinates succinate metabolism. Quantifying all subunits of CII is necessary to confirm this idea though. Increasing SDH/CII flux may be a compensatory mechanism in response to OVX as loss of estrogen has been shown to decrease CI respiration in skeletal muscle^50^. However, this finding contradicts other studies which demonstrate reduced SDH following ovariectomy or ERR KO^51–53^. Most likely, our conflicting data is related to either the protocol employed to prevent estrogens’ effect on skeletal muscle (i.e., reduced estrogen levels versus KO of the receptor) or the length of time between OVX and testing as estrogen levels can fluctuate due to extragonadal aromatisation of estrogen^49^.

Given that OVX altered the concentration of TCA cycle intermediates, we anticipated that proteins associated with metabolism would be impacted by OVX in skeletal muscle. To the contrary, only Rbm20 met the statistically significant criteria (0.75 log_2_ fold change, adjusted *p*<0.05), which was downregulated following OVX. To our knowledge, this is the first time that the loss OVX has been shown to impact Rbm20 expression. Rbm20 is a post-transcriptional splicer abundant in both skeletal and cardiac muscle and known to have more than 30 splicing targets (for a comprehensive review, see^43^). These targets have functions in various pathways, including cytoskeleton organisation^54,55^, contraction^54^ and calcium handling^54,55^, all of which are affected in DMD^56,57^. More recently, loss of Rbm20 has been linked to impaired mitochondrial function and dysregulated metabolic pathways and processes. In Rbm20 KO rats, mitochondrial respiration, particularly driven by CI, is reduced^58^ and metabolic pathways are downregulated^59^, indicating that Rbm20 is an essential regulator of metabolism. While our pathways analysis of the proteome did not indicate up- or downregulation of metabolic pathways (OVX vs Sham), E2 replacement did modulate many metabolic and mitochondrial processes, which were associated with upregulation of Rbm20, strengthening the idea that Rbm20 protein expression is responsive to fluctuations in estrogen. Further investigations are required to better understand this relationship.

E2 replacement upregulated various metabolic and mitochondrial pathways suggesting proteins associated with mitochondrial dynamics, the TCA cycle and the ETC could be affected. While most of the proteins investigated were not statistically different following OVX (compared to Sham), our metabolomics data suggested changes in mitochondrial dynamics and/or TCA function, indicating biologically relevant (albeit non-statistically significant) changes to the mitochondrial proteome. The loss of estrogen/OVX or ERRα KO downregulates metabolism^48,51–53,60^ and our data infer early adaptations at 4-5 weeks post-OVX. Increasing the duration of estrogen deprivation would likely lead to widespread reductions in the expression of mitochondrial and metabolic proteins. A similar scenario is observed with E2 replacement, with our analysis suggesting an overall upregulation of mitochondrial dynamic markers and metabolic proteins. For example, upregulation of Mfn1 is associated with improved bioenergetics, calcium handling and excitation-contraction coupling in muscle^61^. Corresponding upregulation of the master mitochondrial DNA transcription factor, Tfam, and transcription of oxidative metabolic machinery (e.g., Ndufaf2, Uqcr11, Cox20, Atpaf2) to facilitate bioenergetic adaptations were observed. Others have also demonstrated that E2 replacement via implant, as used in this study, increases mitochondrial enzyme activity^62,63^. When considered in conjunction with our data, this suggests that estrogen controls both enzyme expression and kinetics via orchestrating substrate flux. It is important to note however, that implantable hormone systems deliver sustained E2 concentrations in comparison to the cyclical circadian release that is observed *in vivo*^60^, which could lead to supraphysiological levels^62^ and evoke an exaggerated mitochondrial/metabolic response.

Pc was the second most upregulated protein following E2 replacement indicating this could be a primary mechanism of estrogen’s effect on metabolism. OVX downregulated Pc (log_2_ fold change =-0.561, *p*<0.001) but had no impact on any of the other proteins associated with pyruvate metabolism. Pc is responsible for the conversion of pyruvate to oxaloacetate, a TCA cycle intermediate, and its anaplerotic action is critical for pacing TCA cycle flux while fostering biosynthesis pathways (e.g., gluconeogenesis and lipogenesis^64^). Pc KO mice have impaired TCA cycle metabolism, resulting in a ∼50% reduction in TCA cycle intermediates^65^. While pyruvate can also be metabolized by a second enzyme, pyruvate dehydrogenase (PDH), which converts pyruvate into acetyl CoA to drive the TCA cycle, we did not observe any differences in the PDH subunits following OVX nor the pyruvate transporters (Mpc1 and Mpc2). Our data support previous findings indicating that OVX-induced reductions in substrate metabolism are not due to changes in the protein expression of PDH, Mcp1 or Mcp2^60^. Instead, our data suggest that the downregulation of Pc may be partially responsible for the reduction in TCA cycle intermediates, particularly since E2 replacement upregulated Pc expression and restored the concentration of TCA cycle intermediates.

Dilated cardiomyopathy is a common complication of female DMD carriers^66–70^ that is often attributed to the mosaic dystrophin expression observed in cardiac tissue^66,67,71^. Our data indicate that reduced Rbm20 may also contribute to this morbidity, particularly in peri-menopausal female carriers given OVX downregulated Rbm20 expression. Loss of function mutation of Rbm20 causes dilated cardiomyopathy (both of genetic and non-genetic origins^43^). Rbm20 KO rodents and human iPSC-derived Rbm20 mutant cardiomyocytes show impaired cardiac contractility and myocardial stiffness, which is attributed to mis-splicing of calcium handling genes e.g., *Ryr2* and *Camk2d*^72,73^, as well as impaired Frank-Starling mechanism due to mis-splicing of *TTN* (titin^54,74^). These same pathophysiological features were recently documented in human iPSC-derived cardiomyocytes from a female Becker MD carrier engineered to have a DMD mutation^75^. While neither Rbm20 expression, nor its splicing targets, were investigated in the study by Kameda *et al.*, the similarities in cardiac abnormalities between Rbm20-induced dilated cardiomyopathy and cardiomyopathy in female DMD carriers are evident and suggest that further research is warranted to ascertain the role of Rbm20 in female DMD carrier cardiomyopathy.

Research on Rbm20 in the context of skeletal muscle is limited. However, evidence suggests that it controls sarcomere assembly and passive elasticity. Loss of Rbm20 shifts Ttn from the stiff to the compliant isoform^76,77^, a phenotype that reduces contractility manifesting as muscle weakness and exercise intolerance^78^. While Ttn isoforms (and levels) have yet to be quantitated in female DMD carriers, muscle weakness and exercise intolerance are common physiological symptoms^79,80^ – further research is warranted to confirm whether Ttn, and indeed Rbm20, are mediators. No effect of OVX or E2 replacement on Ttn protein expression (log_2_ fold=0.0112, *p*=0.901 and log_2_ fold=-0.0847, *p*=0.332, respectively) was observed, but this does not rule out a potential shift in Ttn isoforms. Thyroid hormone was shown to regulate titin isoform transition via Rbm20 in cardiomyocytes^81^, suggesting other hormones, e.g., E2, may also be influential. Furthermore, loss of Rbm20 in skeletal muscle diminishes exercise capacity^82^, which may partly explain the decreased exercise capacity observed in OVX mice.

Recently, Rbm20 was shown to regulate Ttn splicing in skeletal muscle under the control of the muscle circadian clock regulatory genes, *Bmal1* and *Clock*^32^. There is well-established reciprocal regulation between circadian and estrogen signaling (for a review, see Alvord *et al.*, 2022^83^). The suprachiasmatic nucleus (SCN) is the master timekeeper of the circadian rhythm, and although there is no evidence that it is directly modulated by circulating estrogen rhythms, our pathways analysis suggests OVX might alter SCN-mediated timekeeping via changes to retinal function or vice versa. Four vision-related pathways involving a total of 26 modulated proteins were upregulated including retinoid metabolism and transport, visual phototransduction, diseases associated with visual transduction and canonical retinoid cycle in rods (twilight vision). The SCN receives light input from the retina, and via a complex hormonal and neural milieu, orchestrates tissues-specific transcription of Cryptochrome ((*Cry*) *1* and *2*) and Period ((*Per*)*1*, *2* and *3*) genes by *Bmal1* and *Clock*, to induce local circadian effects^84^. ERα and -β are outputs of the molecular clock transcription program but can also directly regulate the estrogen response element (ERE) on specific circadian genes such as *Per2*^84–86^ and *Clock*^87,88^. ERα expression (log_2_ fold=-0.532, *p*=0.13) was not significantly affected by OVX in our muscle proteome but E2 replacement did upregulate expression (log_2_ fold=0.861, *p*=0.0155; ERβ was not detected). In rodent metabolic tissues, OVX results in a significant circadian phase shift of Per1 expression (liver) and/or a rapid decline in circadian phase synchrony (liver and white adipose tissue) with impact on lipid metabolism, insulin sensitivity and glucose tolerance in these tissues. Our data is the first to indicate that estrogen (and lack thereof) regulates molecular clock control of metabolism in skeletal muscle, another highly metabolic tissue, and that Rbm20 is involved in these adaptations.

## 5. Conclusion

To our knowledge, this is the first study to demonstrate a link between the loss of estrogen and changes to the muscle metabolism that may be regulated via Rbm20 and the circadian clock. Our data indicate that estrogen may play a role in reducing the dystrophinopathic characteristics in female *mdx* mice and protects against exercise intolerance. Importantly, OVX downregulated Rbm20 protein expression, which may contribute to common female DMD carrier manifestations (e.g., dilated cardiomyopathy, exercise intolerance). Further research is required to confirm whether the decrease in Rbm20 protein is also associated with a reduction in Rbm20 splicing capacity. While the use of tamoxifen, a selective ERα agonist, does not appear to be clinically beneficial for DMD patients^30^, our data indicates that E2 replacement could be valuable for female DMD carriers (above menopausal age). It does need to be considered, however, that dystrophic-like features in female carriers can manifest from as young as 3 years of age^89^, which indicates that E2 therapy may not be a universal treatment option for carriers. Follow-up studies are warranted to understand whether estrogen levels correlate with severity/onset of dystrophinopathy in female carriers and if E2 replacement is a viable therapeutic pathway.

## Data Availability

Data will be made available upon reasonable request from the corresponding author.

## Author Contributions

A. Lindsay and E. Rybalka conceived and designed the research; C.A. Timpani, S. Kourakis, D. Debrincat, R. Boyer, L.E. Formosa, J.R. Steele, H. Zhang, R.B. Schittenhelm, E. Rybalka and A. Lindsay conducted the experiments and interpreted the results; C.A. Timpani and E. Rybalka prepared the manuscript, and all authors were involved in critically reviewing the manuscript.

## Acknowledgements

This research was supported by a Deakin University HAtCH Grant (awarded to A.L.) and an Institute for Physical Activity and Nutrition Seed Grant (awarded to A.L.). A.L. was supported by a Philip Wrightson Fellowship (Neurological Foundation) and is currently supported by a Sir Charles Hercus Health Research Fellowship (Health Research Council). L.E.F. is supported by a National Health and Medical Research Council (NHMRC) Investigator Grant (GNT2010149) and funding from the Mito Foundation. C.A.T and E.R. are supported by AFM Telethon (France).

## References

1 Timpani, C. A., Hayes, A. & Rybalka, E. Revisiting the dystrophin-ATP connection: How half a century of research still implicates mitochondrial dysfunction in Duchenne Muscular Dystrophy aetiology. Medical hypotheses 85, 1021–1033 (2015).

2 Budzinska, M., Zimna, A. & Kurpisz, M. The role of mitochondria in Duchenne muscular dystrophy. J Physiol Pharmacol 72, doi:10.26402/jpp.2021.2.01 (2021).

3 Lindsay, A., Chamberlain, C. M., Witthuhn, B. A., Lowe, D. A. & Ervasti, J. M. Dystrophinopathy-associated dysfunction of Krebs cycle metabolism. Human Molecular Genetics 28, 942–951, doi:10.1093/hmg/ddy404 (2018).

4 Tsonaka, R. et al. Longitudinal metabolomic analysis of plasma enables modeling disease progression in Duchenne muscular dystrophy mouse models. Human Molecular Genetics 29, 745–755, doi:10.1093/hmg/ddz309 (2020).

5 Dabaj, I. et al. Muscle metabolic remodelling patterns in Duchenne muscular dystrophy revealed by ultra-high-resolution mass spectrometry imaging. Scientific Reports 11, 1906, doi:10.1038/s41598-021-81090-1 (2021).

6 Ramos, S. V., Hughes, M. C., Delfinis, L. J., Bellissimo, C. A. & Perry, C. G. R. Mitochondrial bioenergetic dysfunction in the D2.mdx model of Duchenne muscular dystrophy is associated with microtubule disorganization in skeletal muscle. PLoS One 15, e0237138, doi:10.1371/journal.pone.0237138 (2020).

7 Timpani, C. A. et al. Attempting to compensate for reduced neuronal nitric oxide synthase protein with nitrate supplementation cannot overcome metabolic dysfunction but rather has detrimental effects in dystrophin-deficient mdx muscle. Neurotherapeutics 14, 429–446 (2017).

8 Mareedu, S., Million, E. D., Duan, D. & Babu, G. J. Abnormal Calcium Handling in Duchenne Muscular Dystrophy: Mechanisms and Potential Therapies. Frontiers in Physiology 12, doi:10.3389/fphys.2021.647010 (2021).

9 Rybalka, E., Timpani, C. A., Stathis, C. G., Hayes, A. & Cooke, M. B. Metabogenic and nutriceutical approaches to address energy dysregulation and skeletal muscle wasting in duchenne muscular dystrophy. Nutrients 7, 9734–9767 (2015).

10 Vu Hong, A., Sanson, M., Richard, I. & Israeli, D. A revised model for mitochondrial dysfunction in Duchenne muscular dystrophy. European Journal of Translational Myology 31, doi:10.4081/ejtm.2021.10012 (2021).

11 Luan, P. et al. Urolithin A improves muscle function by inducing mitophagy in muscular dystrophy. Sci Transl Med 13, doi:10.1126/scitranslmed.abb0319 (2021).

12 Onopiuk, M. et al. Mutation in dystrophin-encoding gene affects energy metabolism in mouse myoblasts. Biochemical and Biophysical Research Communications 386, 463–466 (2009).

13 Moore, T. M. et al. Mitochondrial Dysfunction Is an Early Consequence of Partial or Complete Dystrophin Loss in mdx Mice. Frontiers in Physiology 11, doi:10.3389/fphys.2020.00690 (2020).

14 Mohiuddin, M. et al. Transplantation of Muscle Stem Cell Mitochondria Rejuvenates the Bioenergetic Function of Dystrophic Muscle. bioRxiv, 2020.2004.2017.017822, doi:10.1101/2020.04.17.017822 (2020).

15 Licandro, S. A. et al. The pan HDAC inhibitor Givinostat improves muscle function and histological parameters in two Duchenne muscular dystrophy murine models expressing different haplotypes of the LTBP4 gene. Skelet Muscle 11, 19, doi:10.1186/s13395-021-00273-6 (2021).

16 Timpani, C. A. et al. Dimethyl fumarate modulates the dystrophic disease program following short-term treatment. JCI Insight 8, doi:10.1172/jci.insight.165974 (2023).

17 Lee, T. et al. Differences in carrier frequency between mothers of Duchenne and Becker muscular dystrophy patients. J Hum Genet 59, 46–50, doi:10.1038/jhg.2013.119 (2014).

18 Barbiroli, B., Funicello, R., Ferlini, A., Montagna, P. & Zaniol, P. Muscle energy metabolism in female DMD/BMD carriers: A 31P-MR spectroscopy study. Muscle & Nerve 15, 344–348, 10.1002/mus.880150313 (1992).

19 Barbiroli, B. et al. Further impairment of muscle phosphate kinetics by lengthening exercise in DMD/BMD carriers: An in vivo 31P-NMR spectroscopy study. Journal of the Neurological Sciences 119, 65–73, 10.1016/0022-510X(93)90192-2 (1993).

20 Gaines, R. F., Pueschel, S. M., Sassaman, E. A. & Driscoll, J. L. Effect of exercise on serum creatine kinase in carriers of Duchenne muscular dystrophy. Journal of Medical Genetics 19, 4–7, doi:10.1136/jmg.19.1.4 (1982).

21 Giliberto, F. et al. Symptomatic female carriers of Duchenne muscular dystrophy (DMD): Genetic and clinical characterization. Journal of the Neurological Sciences 336, 36–41, 10.1016/j.jns.2013.09.036 (2014).

22 Song, T. J., Lee, K. A., Kang, S. W., Cho, H. & Choi, Y. C. Three cases of manifesting female carriers in patients with Duchenne muscular dystrophy. Yonsei Med J 52, 192–195, doi:10.3349/ymj.2011.52.1.192 (2011).

23 Quak, Z. X. et al. A manifesting female carrier of Duchenne muscular dystrophy: Importance of genetics for the dystrophinopathies. Singapore Med J 64, 81–87, doi:10.4103/singaporemedj.SMJ-2021-356 (2023).

24 Ohtani, H., Saotome, M., Sakamoto, A., Suwa, K. & Maekawa, Y. Drug-refractory Heart Failure in Female Carrier of Duchenne Muscular Dystrophy: A Case of X-linked Dilated Cardiomyopathy. Intern Med 62, 2089–2092, doi:10.2169/internalmedicine.0745-22 (2023).

25 Lindsay, A. & Russell, A. P. The unconditioned fear response in dystrophin-deficient mice is associated with adrenal and vascular function. Scientific Reports 13, 5513 (2023).

26 Lindsay, A. et al. Sensitivity to behavioral stress impacts disease pathogenesis in dystrophin□deficient mice. The FASEB Journal 35, e22034 (2021).

27 Lowe, D. A. & Kararigas, G. Editorial: New Insights into Estrogen/Estrogen Receptor Effects in the Cardiac and Skeletal Muscle. Frontiers in Endocrinology 11, doi:10.3389/fendo.2020.00141 (2020).

28 Dorchies, O. M. et al. The Anticancer Drug Tamoxifen Counteracts the Pathology in a Mouse Model of Duchenne Muscular Dystrophy. The American Journal of Pathology 182, 485–504, 10.1016/j.ajpath.2012.10.018 (2013).

29 Birnbaum, F., Eguchi, A., Pardon, G., Chang, A. C. Y. & Blau, H. M. Tamoxifen treatment ameliorates contractile dysfunction of Duchenne muscular dystrophy stem cell-derived cardiomyocytes on bioengineered substrates. npj Regenerative Medicine 7, 19, doi:10.1038/s41536-022-00214-x (2022).

30 Henzi, B. C. et al. Safety and efficacy of tamoxifen in boys with Duchenne muscular dystrophy (TAMDMD): a multicentre, randomised, double-blind, placebo-controlled, phase 3 trial. The Lancet Neurology 22, 890–899, doi:10.1016/S1474-4422(23)00285-5 (2023).

31 Nagy, S. et al. Tamoxifen in Duchenne muscular dystrophy (TAMDMD): study protocol for a multicenter, randomized, placebo-controlled, double-blind phase 3 trial. Trials 20, 637, doi:10.1186/s13063-019-3740-6 (2019).

32 Riley, L. A. et al. The skeletal muscle circadian clock regulates titin splicing through RBM20. eLife 11, e76478, doi:10.7554/eLife.76478 (2022).

33 Timpani, C. A. et al. Adenylosuccinic acid therapy ameliorates murine Duchenne Muscular Dystrophy. Sci Rep 10, 1125, doi:10.1038/s41598-020-57610-w (2020).

34 Rybalka, E., Timpani, C. A., Cooke, M. B., Williams, A. D. & Hayes, A. Defects in mitochondrial ATP synthesis in dystrophin-deficient mdx skeletal muscles may be caused by complex I insufficiency. PloS one 9 (2014).

35 HaileMariam, M. et al. S-Trap, an Ultrafast Sample-Preparation Approach for Shotgun Proteomics. Journal of Proteome Research 17, 2917–2924, doi:10.1021/acs.jproteome.8b00505 (2018).

36 Shah, A. D., Goode, R. J. A., Huang, C., Powell, D. R. & Schittenhelm, R. B. LFQ-Analyst: An Easy-To-Use Interactive Web Platform To Analyze and Visualize Label-Free Proteomics Data Preprocessed with MaxQuant. Journal of Proteome Research 19, 204–211, doi:10.1021/acs.jproteome.9b00496 (2020).

37 Zhang, H. et al. Phospho-Analyst: An Interactive, Easy-to-Use Web Platform To Analyze Quantitative Phosphoproteomics Data. Journal of Proteome Research 22, 2890–2899, doi:10.1021/acs.jproteome.3c00186 (2023).

38 Zhang, X. et al. Proteome-wide identification of ubiquitin interactions using UbIA-MS. Nat Protoc 13, 530–550, doi:10.1038/nprot.2017.147 (2018).

39 Ritchie, M. E. et al. limma powers differential expression analyses for RNA-sequencing and microarray studies. Nucleic Acids Research 43, e47–e47, doi:10.1093/nar/gkv007 (2015).

40 Gillespie, M. et al. The reactome pathway knowledgebase 2022. Nucleic Acids Research 50, D687–D692, doi:10.1093/nar/gkab1028 (2021).

41 Dutta, T. et al. Concordance of Changes in Metabolic Pathways Based on Plasma Metabolomics and Skeletal Muscle Transcriptomics in Type 1 Diabetes. Diabetes 61, 1004–1016, doi:10.2337/db11-0874 (2012).

42 O’Leary, M. F., Jackman, S. R. & Bowtell, J. L. Shatavari supplementation in postmenopausal women alters the skeletal muscle proteome and pathways involved in training adaptation. European Journal of Nutrition, doi:10.1007/s00394-023-03310-w (2024).

43 Koelemen, J., Gotthardt, M., Steinmetz, L. M. & Meder, B. RBM20-Related Cardiomyopathy: Current Understanding and Future Options. J Clin Med 10, doi:10.3390/jcm10184101 (2021).

44 Cotta, A. et al. Phenotypic Variability of Dystrophinopathy Symptomatic Female Carriers. Canadian Journal of Neurological Sciences 44, 304–310, doi:10.1017/cjn.2016.448 (2017).

45 Romero, N. B. et al. Pseudo-metabolic presentation in a Duchenne muscular dystrophy symptomatic carrier with ‘de novo’ duplication of dystrophin gene. Neuromuscular Disorders 11, 494–498, 10.1016/S0960-8966(01)00192-4 (2001).

46 Ikeda, K., Horie-Inoue, K. & Inoue, S. Functions of estrogen and estrogen receptor signaling on skeletal muscle. The Journal of Steroid Biochemistry and Molecular Biology 191, 105375, 10.1016/j.jsbmb.2019.105375 (2019).

47 Pellegrino, A., Tiidus, P. M. & Vandenboom, R. Mechanisms of Estrogen Influence on Skeletal Muscle: Mass, Regeneration, and Mitochondrial Function. Sports Medicine 52, 2853–2869, doi:10.1007/s40279-022-01733-9 (2022).

48 Perry, M. C., Dufour, C. R., Tam, I. S., B’Chir, W. & Giguère, V. Estrogen-related receptor-α coordinates transcriptional programs essential for exercise tolerance and muscle fitness. Mol Endocrinol 28, 2060–2071, doi:10.1210/me.2014-1281 (2014).

49 Zhao, H., Tian, Z., Hao, J. & Chen, B. Extragonadal aromatization increases with time after ovariectomy in rats. Reproductive Biology and Endocrinology 3, 6, doi:10.1186/1477-7827-3-6 (2005).

50 Torres, M. J., Ryan, T. E., Lin, C. T., Zeczycki, T. N. & Neufer, P. D. Impact of 17β-estradiol on complex I kinetics and H(2)O(2) production in liver and skeletal muscle mitochondria. J Biol Chem 293, 16889–16898, doi:10.1074/jbc.RA118.005148 (2018).

51 Siebert, C. et al. Effect of physical exercise on changes in activities of creatine kinase, cytochrome c oxidase and ATP levels caused by ovariectomy. Metabolic Brain Disease 29, 825–835, doi:10.1007/s11011-014-9564-x (2014).

52 Wattez, J.-S. et al. Loss of skeletal muscle estrogen-related receptors leads to severe exercise intolerance. Molecular Metabolism 68, 101670, 10.1016/j.molmet.2023.101670 (2023).

53 Torres, M. J. et al. 17β-Estradiol Directly Lowers Mitochondrial Membrane Microviscosity and Improves Bioenergetic Function in Skeletal Muscle. Cell Metab 27, 167–179.e167, doi:10.1016/j.cmet.2017.10.003 (2018).

54 Guo, W. et al. RBM20, a gene for hereditary cardiomyopathy, regulates titin splicing. Nat Med 18, 766–773, doi:10.1038/nm.2693 (2012).

55 Maatz, H. et al. RNA-binding protein RBM20 represses splicing to orchestrate cardiac pre-mRNA processing. The Journal of Clinical Investigation 124, 3419–3430, doi:10.1172/JCI74523 (2014).

56 Rybalka, E. et al. The Failed Clinical Story of Myostatin Inhibitors against Duchenne Muscular Dystrophy: Exploring the Biology behind the Battle. Cells 9, doi:10.3390/cells9122657 (2020).

57 Kourakis, S. et al. Targeting Nrf2 for the treatment of Duchenne Muscular Dystrophy. Redox Biol 38, 101803, doi:10.1016/j.redox.2020.101803 (2021).

58 Vad, O. B. et al. Loss of Cardiac Splicing Regulator RBM20 Is Associated With Early-Onset Atrial Fibrillation. JACC: Basic to Translational Science, 10.1016/j.jacbts.2023.08.008 (2023).

59 Larson, E. J. et al. Rbm20 ablation is associated with changes in the expression of titin-interacting and metabolic proteins. Molecular Omics 18, 627–634, doi:10.1039/D2MO00115B (2022).

60 Takahashi, K., Kitaoka, Y. U., Matsunaga, Y. & Hatta, H. Effects of Endurance Training on Metabolic Enzyme Activity and Transporter Proteins in Skeletal Muscle of Ovariectomized Mice. Med Sci Sports Exerc 55, 186–198, doi:10.1249/mss.0000000000003045 (2023).

61 Eisner, V., Lenaers, G. & Hajnóczky, G. Mitochondrial fusion is frequent in skeletal muscle and supports excitation-contraction coupling. J Cell Biol 205, 179–195, doi:10.1083/jcb.201312066 (2014).

62 Campbell, S. E. & Febbraio, M. A. Effect of ovarian hormones on mitochondrial enzyme activity in the fat oxidation pathway of skeletal muscle. Am J Physiol Endocrinol Metab 281, E803–808, doi:10.1152/ajpendo.2001.281.4.E803 (2001).

63 Beckett, T., Tchernof, A. & Toth, M. J. Effect of ovariectomy and estradiol replacement on skeletal muscle enzyme activity in female rats. Metabolism 51, 1397–1401, doi:10.1053/meta.2002.35592 (2002).

64 Jitrapakdee, S. et al. Structure, mechanism and regulation of pyruvate carboxylase. Biochem J 413, 369–387, doi:10.1042/bj20080709 (2008).

65 Cappel, D. A. et al. Pyruvate-Carboxylase-Mediated Anaplerosis Promotes Antioxidant Capacity by Sustaining TCA Cycle and Redox Metabolism in Liver. Cell Metab 29, 1291–1305.e1298, doi:10.1016/j.cmet.2019.03.014 (2019).

66 Adachi, K. et al. Detection and management of cardiomyopathy in female dystrophinopathy carriers. Journal of the Neurological Sciences 386, 74–80, 10.1016/j.jns.2017.12.024 (2018).

67 Politano, L. et al. Development of Cardiomyopathy in Female Carriers of Duchenne and Becker Muscular Dystrophies. JAMA 275, 1335–1338, doi:10.1001/jama.1996.03530410049032 (1996).

68 Hoogerwaard, E. M. et al. Signs and symptoms of Duchenne muscular dystrophy and Becker muscular dystrophy among carriers in The Netherlands: a cohort study. Lancet 353, 2116–2119, doi:10.1016/s0140-6736(98)10028-4 (1999).

69 Grain, L. et al. Cardiac abnormalities and skeletal muscle weakness in carriers of Duchenne and Becker muscular dystrophies and controls. Neuromuscul Disord 11, 186–191, doi:10.1016/s0960-8966(00)00185-1 (2001).

70 Schelhorn, J. et al. Cardiac pathologies in female carriers of Duchenne muscular dystrophy assessed by cardiovascular magnetic resonance imaging. European Radiology 25, 3066–3072, doi:10.1007/s00330-015-3694-3 (2015).

71 Florian, A. et al. Cause of Cardiac Disease in a Female Carrier of Duchenne Muscular Dystrophy. Circulation 129, e482–e484, doi:doi:10.1161/CIRCULATIONAHA.113.006891 (2014).

72 Guo, W. et al. Splicing Factor RBM20 Regulates Transcriptional Network of Titin Associated and Calcium Handling Genes in The Heart. Int J Biol Sci 14, 369–380, doi:10.7150/ijbs.24117 (2018).

73 Hoogenhof, M. M. G. v. d., et al. RBM20 Mutations Induce an Arrhythmogenic Dilated Cardiomyopathy Related to Disturbed Calcium Handling. Circulation 138, 1330–1342, doi:doi:10.1161/CIRCULATIONAHA.117.031947 (2018).

74 Beqqali, A. et al. A mutation in the glutamate-rich region of RNA-binding motif protein 20 causes dilated cardiomyopathy through missplicing of titin and impaired Frank-Starling mechanism. Cardiovasc Res 112, 452–463, doi:10.1093/cvr/cvw192 (2016).

75 Kameda, S. et al. Modeling Reduced Contractility and Stiffness Using iPSC-Derived Cardiomyocytes Generated From Female Becker Muscular Dystrophy Carrier. JACC: Basic to Translational Science 8, 599–613, 10.1016/j.jacbts.2022.11.007 (2023).

76 Buck, D. et al. Removal of immunoglobulin-like domains from titin’s spring segment alters titin splicing in mouse skeletal muscle and causes myopathy. Journal of General Physiology 143, 215–230, doi:10.1085/jgp.201311129 (2014).

77 Ma, W., Buck, D., Nedrud, J., Irving, T. & Granzier, H. Thick Filament Compliance in Passively Stretched Skeletal Muscle. Biophysical Journal 112, 181a, doi:10.1016/j.bpj.2016.11.1005 (2017).

78 Pijl, R. J. v. d., Granzier, H. L. & Ottenheijm, C. A. C. Diaphragm contractile weakness due to reduced mechanical loading: role of titin. American Journal of Physiology-Cell Physiology 317, C167–C176, doi:10.1152/ajpcell.00509.2018 (2019).

79 Papa, R. et al. Genetic and Early Clinical Manifestations of Females Heterozygous for Duchenne/Becker Muscular Dystrophy. Pediatric Neurology 55, 58–63, 10.1016/j.pediatrneurol.2015.11.004 (2016).

80 Kakulas, B. A. in Exercise Intolerance and Muscle Contracture (eds Georges Serratrice, Jean Pouget, & Jean-Philippe Azulay) 75–82 (Springer Paris, 1999).

81 Zhu, C. et al. RBM20 is an essential factor for thyroid hormone-regulated titin isoform transition. J Mol Cell Biol 7, 88–90, doi:10.1093/jmcb/mjv002 (2015).

82 Guo, W., Pleitner, J. M., Saupe, K. W. & Greaser, M. L. Pathophysiological Defects and Transcriptional Profiling in the RBM20-/- Rat Model. PLOS ONE 8, e84281, doi:10.1371/journal.pone.0084281 (2013).

83 Alvord, V. M., Kantra, E. J. & Pendergast, J. S. Estrogens and the circadian system. Semin Cell Dev Biol 126, 56–65, doi:10.1016/j.semcdb.2021.04.010 (2022).

84 Nakamura, T. J. et al. Estrogen differentially regulates expression of Per1 and Per2 genes between central and peripheral clocks and between reproductive and nonreproductive tissues in female rats. J Neurosci Res 82, 622–630, doi:10.1002/jnr.20677 (2005).

85 Perrin, J. S., Segall, L. A., Harbour, V. L., Woodside, B. & Amir, S. The expression of the clock protein PER2 in the limbic forebrain is modulated by the estrous cycle. Proc Natl Acad Sci U S A 103, 5591–5596, doi:10.1073/pnas.0601310103 (2006).

86 Nakamura, T. J., Sellix, M. T., Menaker, M. & Block, G. D. Estrogen directly modulates circadian rhythms of PER2 expression in the uterus. Am J Physiol Endocrinol Metab 295, E1025–1031, doi:10.1152/ajpendo.90392.2008 (2008).

87 Xiao, L. et al. Induction of the CLOCK gene by E2-ERα signaling promotes the proliferation of breast cancer cells. PLoS One 9, e95878, doi:10.1371/journal.pone.0095878 (2014).

88 Nakamura, T. J. et al. Influence of the estrous cycle on clock gene expression in reproductive tissues: effects of fluctuating ovarian steroid hormone levels. Steroids 75, 203–212, doi:10.1016/j.steroids.2010.01.007 (2010).

89 Preuße, C. et al. Inflammation-induced fibrosis in skeletal muscle of female carriers of Duchenne muscular dystrophy. Neuromuscular Disorders 29, 487–496, 10.1016/j.nmd.2019.05.003 (2019).

