## Supplementary Figure 1 for "Loss of endogenous estrogen alters mitochondrial metabolism and muscle clock-related protein Rbm20 in female *mdx* mice"

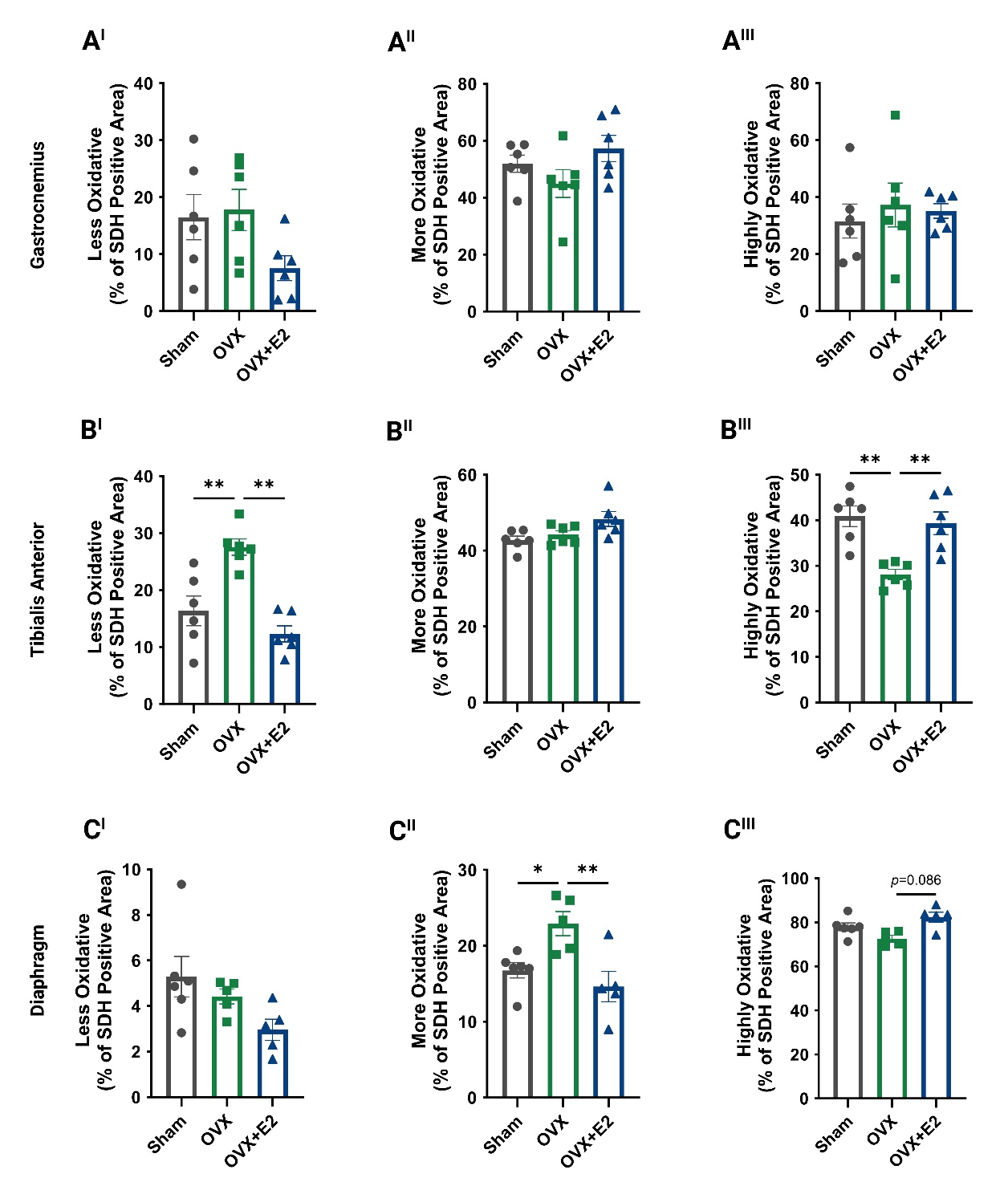

**Supplementary Figure 1. Estrogen depletion has mixed effects on succinate dehydrogenase (SDH) activity in the gastrocnemius, tibialis anterior and diaphragm of female *mdx* mice, which is restored with estradiol (E2).** (A^I-III^) The proportion of less, more, and highly oxidative fibers in gastrocnemius was comparable across the groups. (B^I-III^) In the tibialis anterior, ovariectomy (OVX) drove the proportion of less oxidative fibers to be higher compared to Sham, which was normalized following estrogen replacement while estrogen depletion decreased the proportion of highly oxidative fibers, and this too was normalized with E2 replacement. (C^I-III^) In the diaphragm, OVX increased the number of more oxidative fibers compared to Sham, which was attenuated with E2 replacement. No difference in highly oxidative fibers was observed between Sham and OVX mice, however there was a trend for E2 therapy to increase the proportion of these fibers (*p*=0.086). **p*<0.05, ***p*<0.01. Sham *n*=5; OVX *n*=4; OVX+E2 *n*=5.
